# Physical properties characterization of Natural Protein Fibre Peacock Feather Barbs

**DOI:** 10.1101/238626

**Authors:** Sekhar Das, Seiko Jose, Pintu Pandit

## Abstract

In the present study, the barbs of peacock feather were subjected to its physio-mechanical characterisation. Various properties of barbs viz., bundle strength, diameter, moisture regain, thermal stability, X-ray diffraction, colour intensity and FTIR was studied according standard analytical methods. The surface morphology of the barbs was examined using SEM images. The results indicate that the barb is a hollow vertical structure made up of protein. The average length and diameter of the barb was found to be 45 mm and 82 μm respectively. The FTIR study confirms the presence of characteristic peaks for protein, related to the keratinous material. The barbs seem to be semi-crystalline in nature, as indicated by X-ray study.

## 1. Introduction

Peacock feather attracts people for its beauty, aesthetic appearance and economic value. The feather of a bird performs different important functions such as flight, thermoregulation, swimming, physical protection, decoration, sound production, and foraging and water repellency (Homberger, & de Silva, 2000; Zhang, & Zhou, 2000; Chuong et al., 2000). A feather is a branched structure and composed of a matrix of keratin similar to natural protein fibre hair and fur (Prum, & Williamson, 2001). Feather fibre is build of beta keratin, a unique fibrous protein, in which a filament-matrix structure is formed by each single beta keratin molecule (Greenwold, & Sawyer, 2011; Srinivasan, 2014). As a protein fibre barb has several advantages over commonly available protein fibres like silk and wool. It has low density, unique morphological structure, warmth retention capability, excellent compressibility, resiliency, ability to dampen sound, etc. make them matchless fibers (Barone, & Schmidt, 2005).Feathers are extremely diverse and complex structure in nature (Streit, & Heidrich, 2002). They have complex branched structure and diversity in size, shape, color, and texture. Generally, a feather is made of the calamus that extends into the rachis, the central beam of the feather. The primary branches of the rachis are the barbs and the branches of the barbs are called barbules. Several researchers have worked on the structure and properties of chicken feather barbs and its new application areas. Reddy & Yang (2007) studied the physical and morphological structure and properties of chicken feather barbs and evaluate their suitability as textile fibers. They compared the structure and properties of chicken feather barbs with the protein fiber, wool. Tesfaye et al. (2017) characterized details about the physical properties of a chicken feather. They stated that chicken feather barb has unique feature protein fibre. The barb fibre has low density, high flexibility, good spinning length and a hollow honeycomb structure and suitable for the manufacture of composite materials. In recent years, scientists have chosen biodegradable natural fibre to develop environment-friendly polymer matrix composites (Das et al., 2015; Das et al., 2016; Das et al., 2017a; Das et al., 2017b; Das et al., 2015c;). Several attempts are reported in the literature on using the barbs as “feather fibers” for composites and non-woven applications (Barone, & Schmidt, 2005; Cheng et al., 2009; Kowshik et al., 2017). The peacock feather is attractive and colorful, one of the most well-known examples for the structural color in nature. Several researchers paid great attention to understand the structural color phenomena of a peacock feather and develop novel material inspired by the phenomena. In this regard, Yoshioka, & Kinoshita (2002) studied the structural and reflective properties of peacock feathers. Han et al., (2008) studied optical properties of ZnO nano particles embedded peacock feathers. They claimed that ZnO nano particles embedded peacock feather hybrids would have important applications in optoelectronics and optical communications.

The reported work characterized Contour feather barbs of peacock for their physical and morphological structure and properties. Some physical properties of feather barbs have been compared with the most common natural protein fiber, wool.

## 2. Materials and methods

### 2.1 Sample collection and preparation

India peacocks (Pavo cristatus) normally molt once in a year during the period of July-October. The contour feathers of peacock have collected from ICAR-CSWRI Avikanagar, Rajasthan Campus during these months. No birds were sacrificed specifically for this study. The barbs were separated from the rachis manually by cutting with a sharp blade. The barbs were conditioned at a relative of humidity 65 ± 2 % and a temperature of 20 ± 2°C before testing.

### 2.2 Morphological structure

Scanning electron microscopy (SEM) of the barb is performed with a Philips XL 30 scanning electron microscope.

### 2.3 Length and Diametre

The barb’s length was measured in the interval of 5 cm of rachis length and plotted in the graph. The diameter of 100 barbs was measured using a projection microscope cope. Plots of the barb’s diameter frequency distribution, using the average frequency, in relation to each diameter interval, are shown in Fig.4.

### 2.4 Moisture Regain

Moisture regains was determined according to ASTM D1576-90 standard. Two-gram samples were taken for this purpose barbs were first dried in a hot air oven at 105°C for 4 h. The dried barbs were allowed to regain moisture under the standard testing conditions of 21 °C and 65% RH. The ratio of the dry weight of the barbs to the conditioned weight was taken as the % moisture regain.

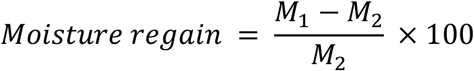

*M*_1_ = Original mass of sample (g), and *M*_2_ = Oven dry mass ofsample (g).

### 2.5 Bundle Strength measurements

Barb fibre bundle strength was determined at standard testing condition in terms of the fibre-bundle tenacity (g/tex) on a Statex Fibre Bundle Strength Tester keeping gauge length zero.

### 2.6 Fourier transforms infrared spectroscopy analysis

The Fourier transform infrared spectroscopy (FTIR) analysis of the barb sample was carried out in a Bruker Alpha-T FTIR spectrometer over the wavelength of 500 to 4000 cm^−1^.

### 2.7 X-ray diffraction analysis

Wide-angle X-ray diffraction (XRD) analysis of the father samples was carried out on Shimadzu 6100, equipped with CuKα radiation (*λ*=1.54 Å) in the 2θ ranging from 5 to 70°. Generator voltage was 40KV, generator current was 30 mA, in step of 0.02°. The sample was prepared as a chopped feather and placed on the stub. The XRD diffraction patterns are presented in Fig. 3. The 2θ values were calculated using the following Eq. (1)

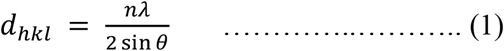

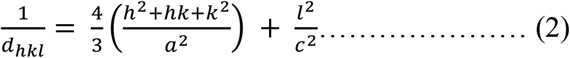

Where θ is the angle of diffraction, n is the wavelength of the X-ray, and dhkl is inter atomic spacing for atoms with Miller indices (hkl), crystallite dimension in the direction perpendicular to the crystallographic plane hkl. Fether exhibited the hexagonal form. For hexagonal crystals, a = b ≠ c and *α* = *β* = 90°; *γ* = 120° where a, b, c and a, β and γ are the lattice parameters. The crystallinity index (C_I_) was measured from the following Eq

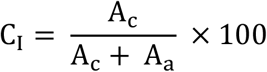

### 2.8 Thermo Gravimetric Analysis

The Thermal Gravimetric Analysis (TGA) of peacock feather samples was carried out using Shimadzu 60H DTG in the temperature range of 30-500°C with a heating rate of 10°C/min under an inert atmosphere of nitrogen at a flow rate of 50 ml/min. The temperature accuracy of the instrument was ±0.3 °C, with a reproducibility of ±0.1 °C; the weighing precision was 1μg, with a sensitivity of 0.1μg, and a dynamic range of ±500mg, having a measurement accuracy of ±1%.

### 2.9 Evaluation of Coloration on father sample

The color depth of the peacock feather sample was evaluated by measuring the reflectance values on a computer color matching system (Spectra scan 5100+ spectrophotometer) at D65 illuminate /10°observer. The Kubelka-Munk function, K/S, which is proportional to the color strength, was determined using the following equation:

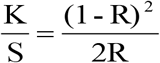

where K is the absorption coefficient, S is the scattering coefficient, and R is the reflectance of the colour at λ max. Other calorimetric value such as L* (lightness - darkness), a* (red - green), and b* (blue - yellow) were also evaluated.

## 3. Results and Discussion

### 3.1 Morphological Structure

The morphological features of contour feather barbs are shown in Fig. 1 & 2. A feather is mainly composed of three different parts; the rachis, the central shaft of the feather that extends the entire length of the feather, the secondary structures, the barbs which are attached to the rachis, the tertiary unit, the barbules are connected to the barbs in a manner similar to the barbs being attached to the rachis. The longitudinal view of barbs surface is smooth and does not contain any scales like protein fibre wool. The cross-section view of barbs is a unique that is not seen in the natural protein fibers wool and silk. The SEM images Fig. confirm that the cross-section of the barb is nearly round or elliptical shape. The inner section of the barbis typically thin wall rectangular to oval shape hollow structure indicate the presence of extensive air pockets in the structure which may be used in the preparation of good thermal retention and light weight materials (Butler, & Johnson, 2004).

**Fig. 1.**
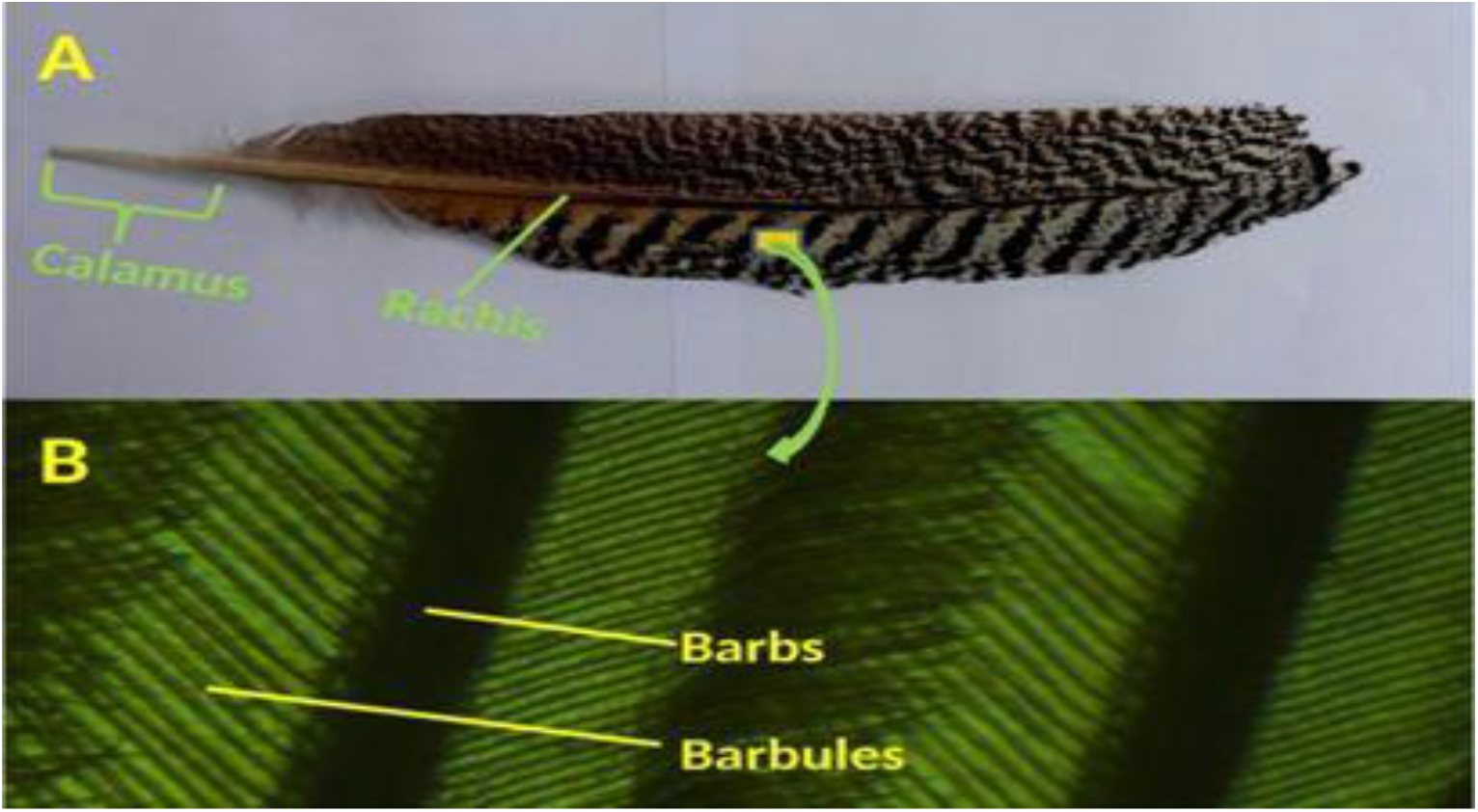
(A) The structure of a typical peacock contour feather (B) Microscopic image of barb and barbules

**Fig. 2.**
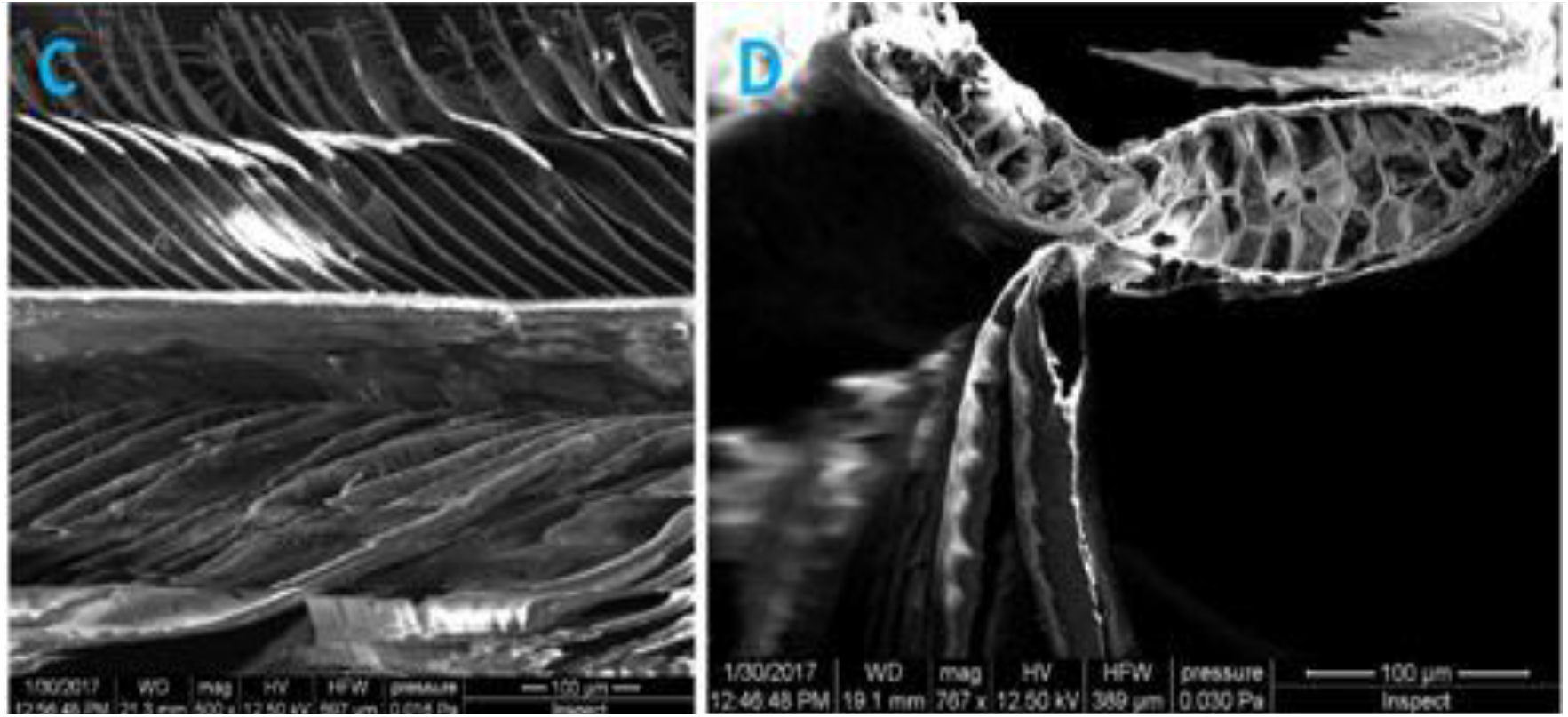
Morphological structures of peacock contour feathers (A) SEM image of the barb surface (B)SEM picture of the cross-section of a barb showing the hollow structures

### 3.2 Barbs Fibre Length

The fig.3 illustrates the barb length variation along the length of rachis of peacock feathers. The length distributions of the feather barb along the length of one rachis were not consistent along their lengths. The fibre length at the base of rachis was around 10 mm and tip fibre length 8 mm of a 48 mm long rachis. The average length of barb was 45 mm with standard deviation 23.13. Peacock feather vane-width asymmetry is mainly caused by barb length. Vane-width asymmetry geometry of feather due to barb length plays significant role aerodynamic performance and portion of the feather vane (Feo, Field, & Prum, 2015).

**Fig. 3.**
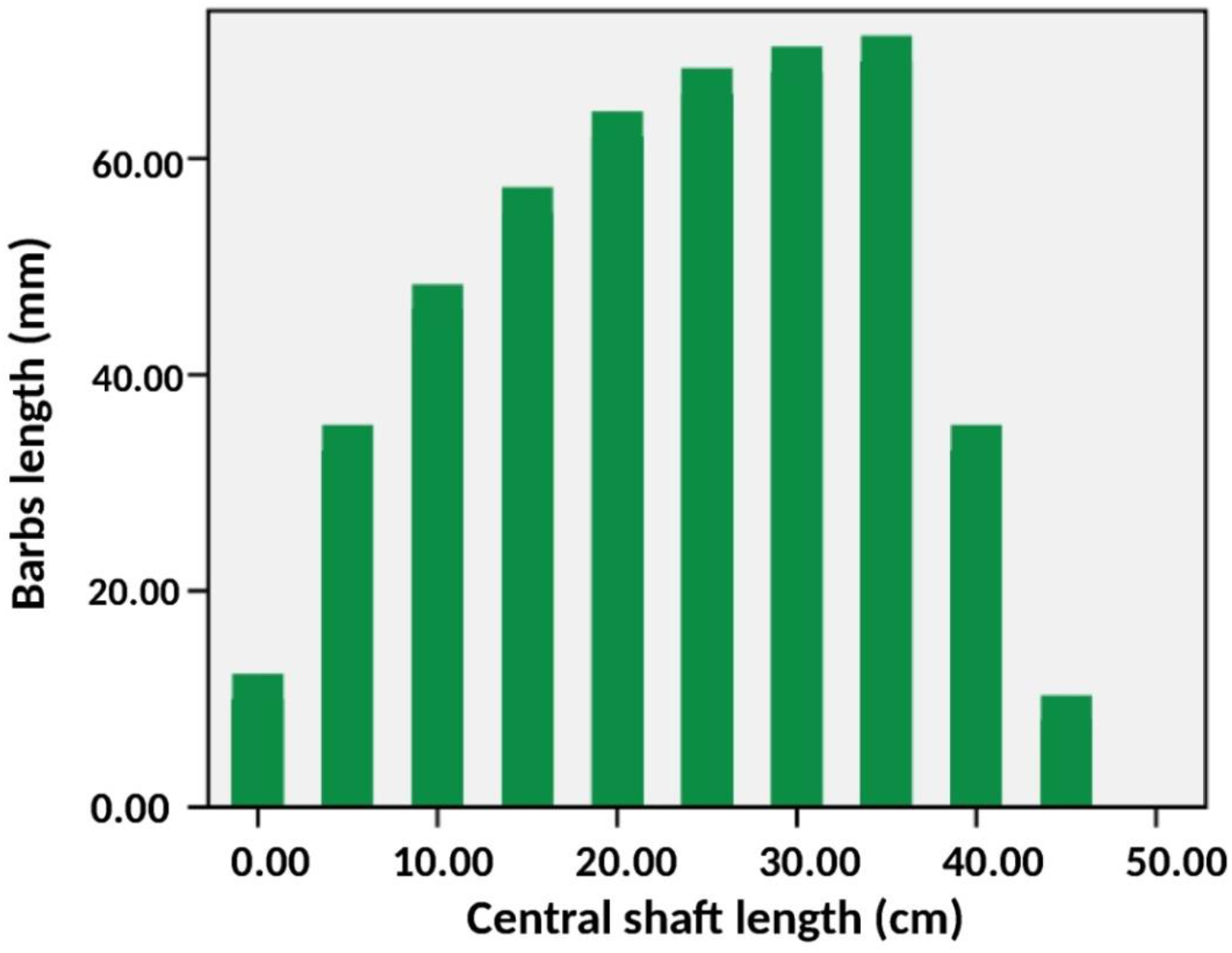
Barb’s length distribution respect to the length of rachis

### 3.3 Barbs Fibre diameter

The fig.4 illustrates the diameter variation of the barbs of peacock feathers. The averages of 100 readings from different places along a single sample were used to calculate the diameter of the barb. It was observed that the mean diameter of the peacock feather bar was 82.2μm, with a SD of 11.7μm.The diameter of the barb which relates to the aspect ratio (fiber length/diameter) is an important parameter affecting the bending properties of the vane. High aspect ratio indicates more flexible barb. The aspect ratio of barb plays significant role aerodynamic performance and portion of the feather.

**Fig. 4.**
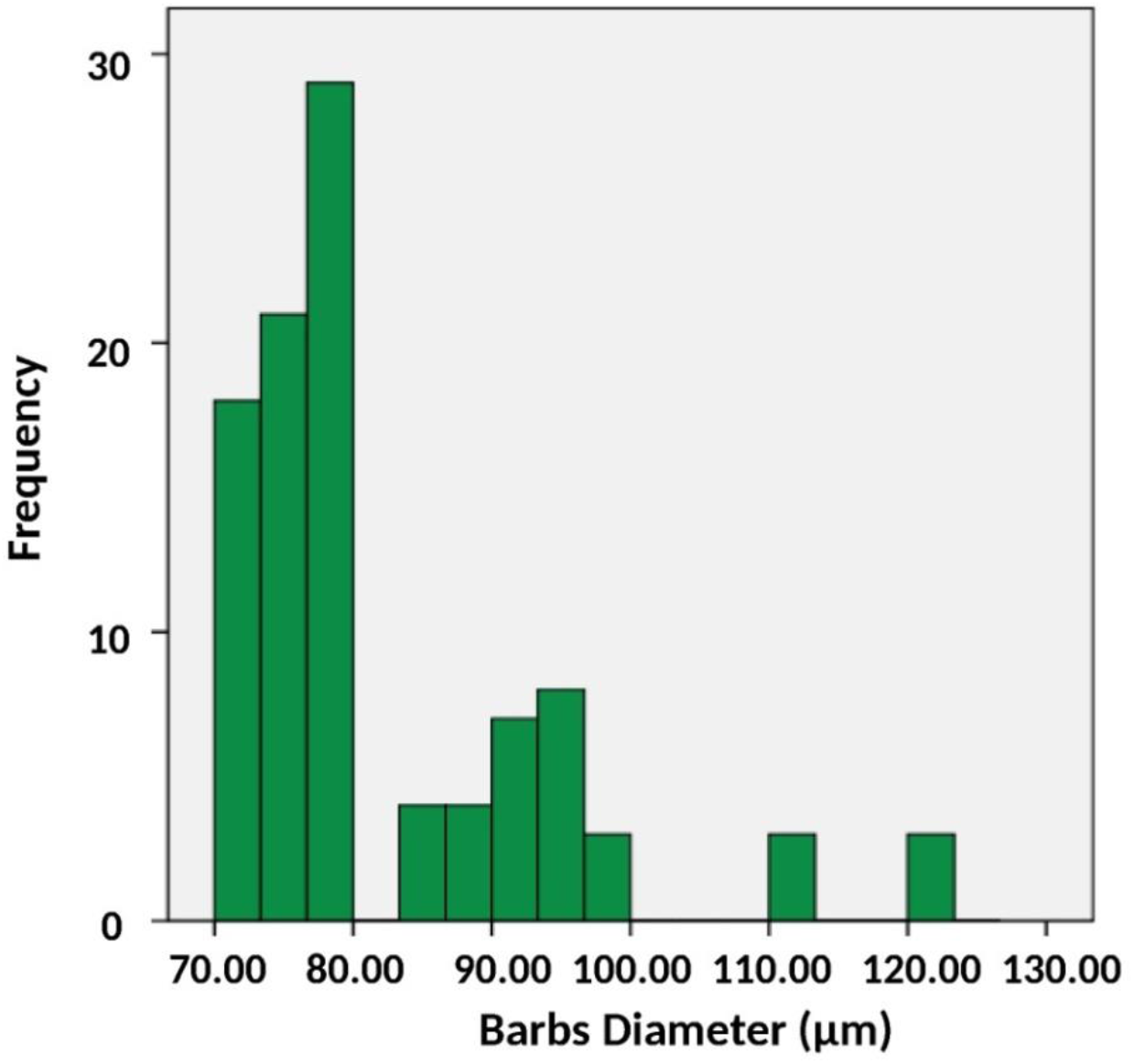
Barb’s diameter frequency distribution

### 3.4 Moisture Regain

It was observed that the moisture regain of feather barbs is 10.6% which is lower than that of protein fibre wool but similar like chicken feather barb (Reddy, & Yang, 2007). The feather fibres are hygroscopic in nature due to presence of polar group which can attach water molecule from atmosphere by creating hydrogen bond (Das, 2017). The amount of moisture that present in barb fiber strongly affects many of their important physical properties such as fibre dimensions; tensile properties, elastic recovery, electrical properties, and thermal properties etc are affected by the amount of water absorbed. It has been observed that when wool fibre absorbs moisture, the initial modulus, yield stress and breaking stress decreases while the breaking extension tends to increase.

### 3.5 Bundle Strength

It was observed that the bundle strength of feather barbs at zero gauge length is 14.84 g/tex with a SD of 1.6. However, zero-span bundle strength results as such tell little of the variation in single fiber strength. Still there is strong correlation between bundle strength and single fibre tensile test. Since the fibre bundle strength is commonly used as a fiber strength index, it also gives information about fiber deformations, fiber damage and variation in individual fiber strand strength. The strength of barbs is highly related to feather’s different functions. The barbs have to be sufficiently strong to withstand aerodynamic forces generated on barbs during flight.

### 3.6 Fourier transforms infrared spectroscopy analysis of barb feather

The fig. 5 shows the FTIR spectrum of the feather presents characteristic bands of protein, relating to the keratinous material. Infrared absorption spectra of feather authenticate characteristic absorption bands assigned mainly to the peptide bonds (–CONH–). The peptide bonds vibrations create bands known as amides I–III (Han, 2008; Sun, 2009). The amide I band is associated mostly with the C=O stretching vibration and it shows in the range of 1700 – 1600 cm^−1^. The peaks 1626 cm^−1^corresponds to elastic vibration of C=O bond. The amide III peaks are shown in the range of 1220–1300 cm^1^. The peak at 1232 cm^−1^ indicates the group CNH in the wool fiber. The amide III band is associated with the C–N stretching, N–H in-plane bending, C–C stretching and C=O bending vibrations. The amide III band of the feather is observed at 1231cm^−1^. In addition, the C–S stretching peaks at 817 cm^−1^ and C¾bending at 1451 cm^−1^ are also observed (Wojciechowska, 1999). The peak at 1525 cm^−1^ is for the bending deformation peak of C–N–H bond (Jose et al., 2018).

**Fig. 5.**
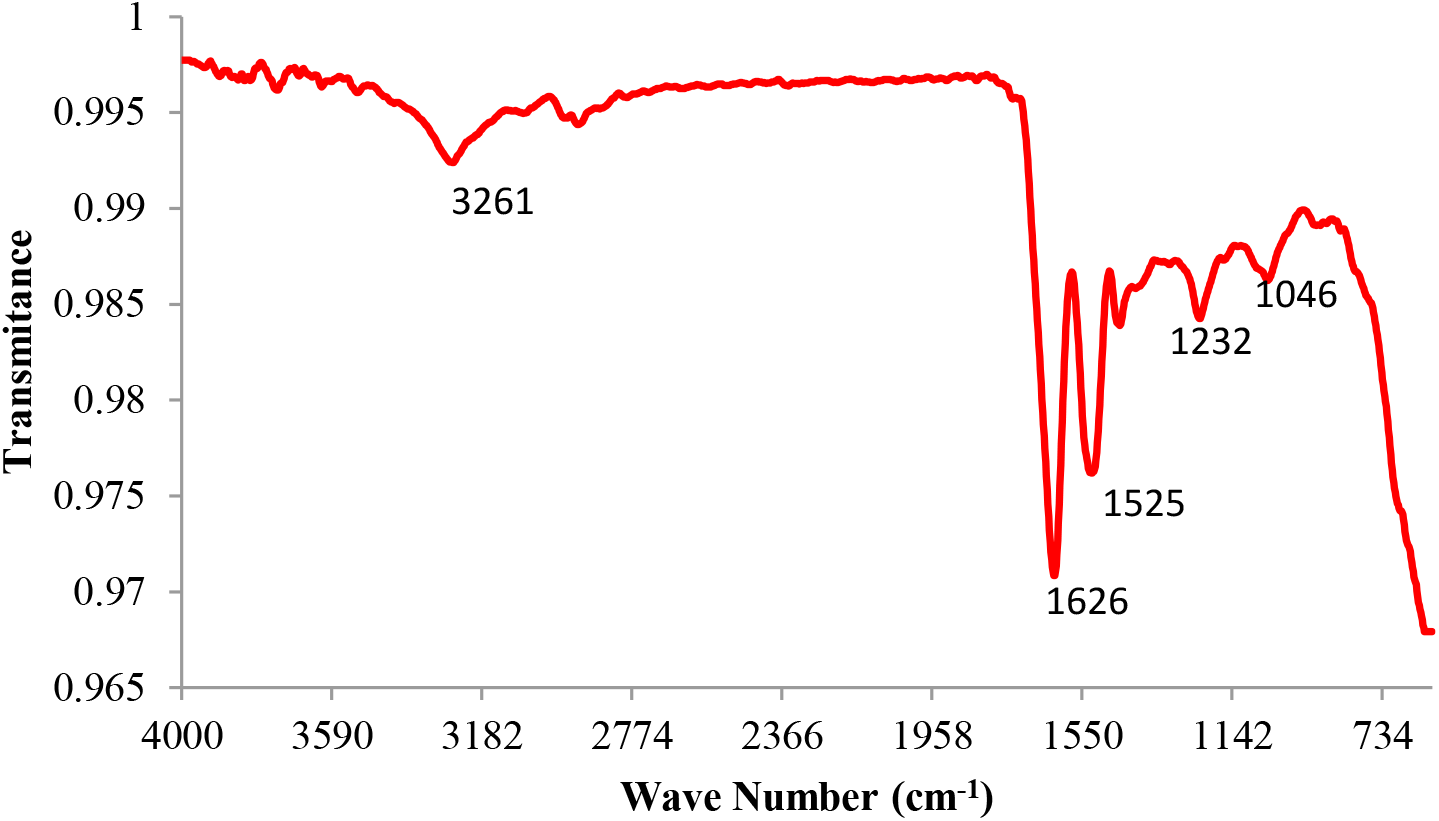
Fourier transforms infrared spectroscopy (FTIR) analysis of peacock feather barb

### 3.7 XRD analysis of barbs fibre

An X-ray diffractogram of feather barb sample is shown in Fig.6 Feather barbs are constructed mainly of beta-keratins, fibrous proteins (Kirschner,1987). The structure of b-type keratin is a twisted b-sheet of laterally packed b-strands and the chains are held together by intermolecular hydrogen bonds (Meyers, 2008; Greenwold, & Sawyer, 2011). The feather barb fibre showed a very broad peak at 9.6° and 19.4° which specifically corresponds to the b-sheet structure and are assigned to [002], and [004], reflections. The equatorial scattering at interplanar distance 9.12 and 4.55 A° demonstrates all of the characteristics of a b-keratin diffraction pattern. The diffraction pattern showed sharp crystallinity peak at 37.6°, 43.9°, 56.7°and 69.9°correspond to the [3-11], [302], [3-10] and [241] plans reflection. It is observed that the feather barb beta keratin unit cell is hexagonal and its dimensions are a = 7.35Å, b = 7.35Å, c = 18.20Å and α=β=90 γ=120. It is reported in the literature that avian beta keratin protein unit cell is orthorhombic and its parameters are a = 9.46 A°, b = 9.7 A° and c = 6.68 (Rizzo, 2006). Further, the crystallinity index of feather barb found to be 15.6%. Feather barb beta keratin fiber is semi-crystalline fibrous material. The crystallinity index play significant role on mechanical properties of fibre. The mechanical properties of the fibre increase with increase in crystallinity index.

**Fig. 6.**
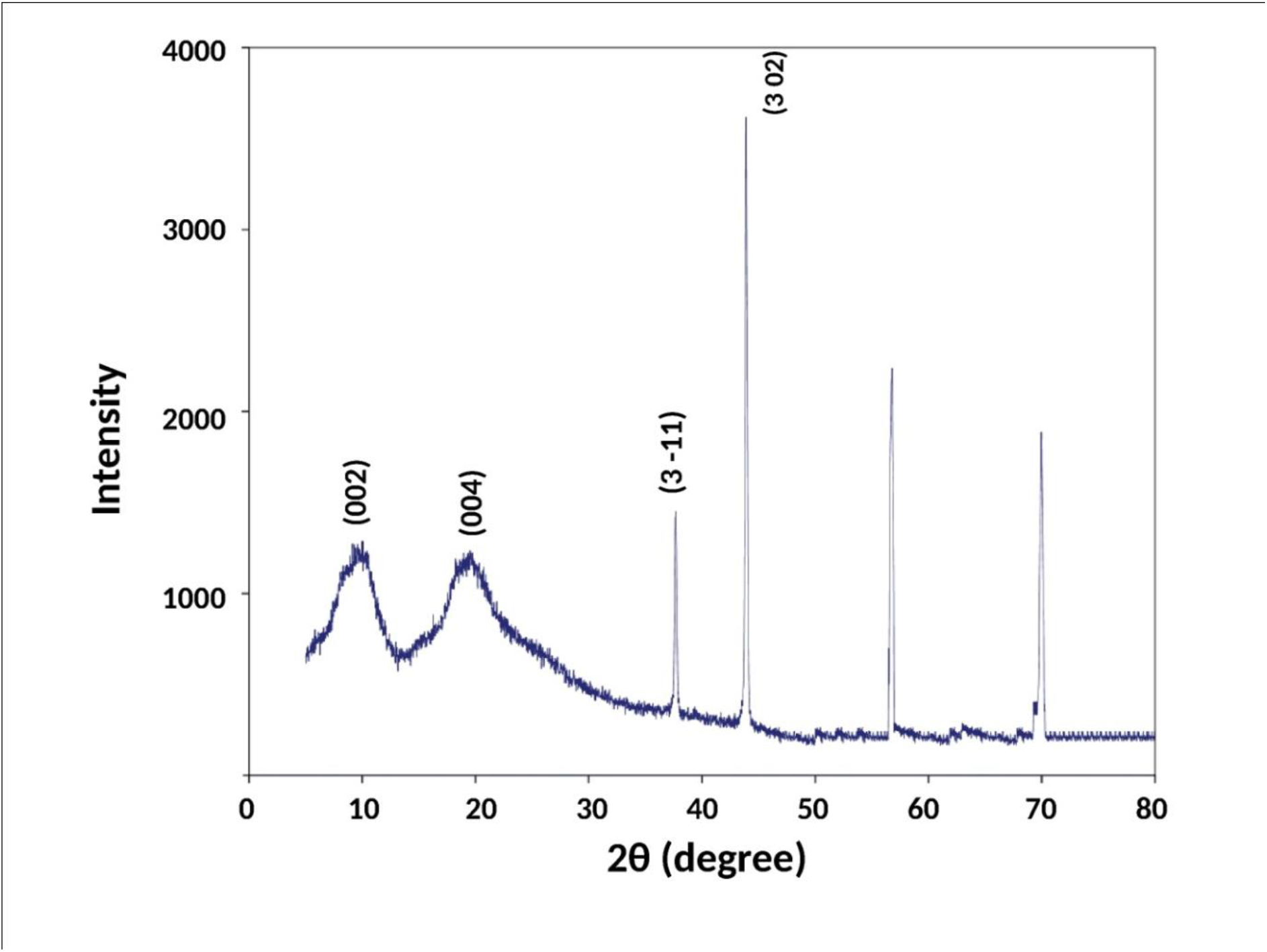
X-ray diffraction spectra of peacock feather barb

### 3.8 Thermogravimetric analysis

The fig.7 shows the thermogravimetric (TG) curves of the feather in the N2 atmosphere at a heating rate of 10°C /min. The TG curve of feather sample shows three main stages of weight loss during the heating process. The initial water desorption process, occurring from 30°C to 150°C and is accompanied by a weight loss of 10%. Three different types of water molecule such as free water, loosely bonded water and chemically bonded are shown to be attached to the feather fiber. The feather fibre has polar amino acid side chains hydrophilic groups and segments suitable for hydrogen bonding which can attach with a water molecule (Senoz, 2012; Cheng, 2009). In the second stage of weight loss process lies from 220°C to 341°C and along with about 21% loss of feather fiber mass, is responsible for the pyrolysis of feather fibre. In the pyrolysis process, the degradation of the protein chain molecules was occurred and produced of sulfur dioxides and hydrogen sulfides due to breakage of disulfide bonds (Khosa,2013; Sharma, 2017). The third region is an exothermic reaction starts from 300°C to 500°C, where weight loss observed to be 43% and the char oxidation reactions are dominated.

**Fig. 7.**
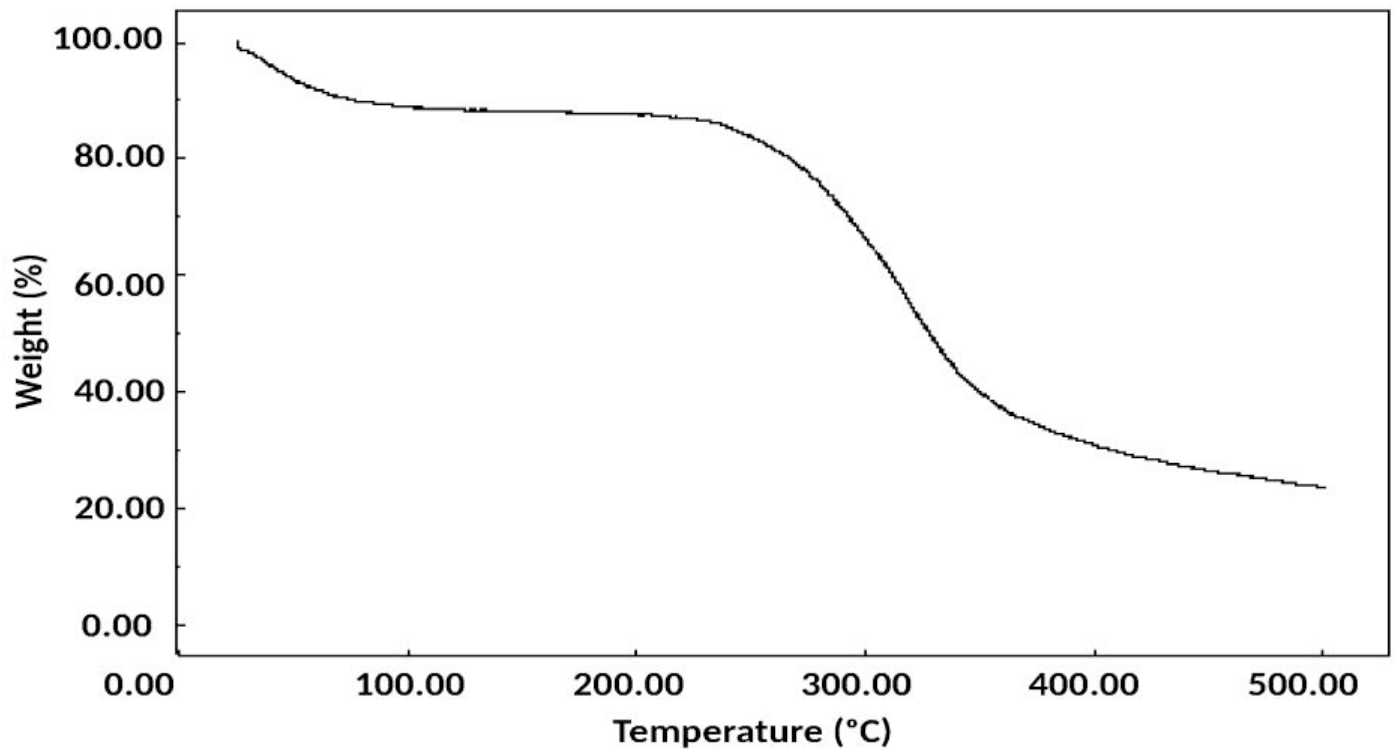
Thermal gravimetric analysis (TGA) curves of peacock feather barb

### 3.9 Colour measurements

The colour strength along with colour co-ordinates L, a*, b*,C*, H* values are reported in Table 1 was analysed using computer colour matching software. The efficacy of feather sample impart coloration in terms of K/S value indicate the total colour value was seen in the case of barb peacock feather. In Table 1, L* values indicate the depth of shade, a* value indicates the tone of the shade in greener or redder region and b* value indicates the tone of the shade in yellow or blue region of the peacock feather.

**Table 1:** Colour co-ordinates values of the peacock feather

| Feather Peacock<br>(at different position) | **L\*** | **a\*** | **b\*** | **C\*** | **H\*** | **K/S** |
| --- | --- | --- | --- | --- | --- | --- |
| 1 | 29.146 | 2.119 | 3.976 | 4.505 | 61.920 | 7.002 |
| 2 | 1.327 | 0.470 | 1.1327 | 1.226 | 0.106 | 10.319 |
| 3 | 4.4041 | 0.707 | 3.382 | 3.377 | 0.732 | 12.739 |
| 4 | 2.041 | -0.326 | 1.856 | 1.596 | 1.002 | 13.146 |
| 5 | 13.531 | -0.671 | 4.173 | 3.771 | 1.908 | 6.035 |

Note: L\*: lightness (0 = black, 100 = white), a\*: red-green coordinates (positive values = red, negative values = green), b\*: yellow-blue coordinates (positive values = yellow, negative values = blue); C\* = Chroma ; H\* = Hue.

## 4. Conclusions

The characterization of peacock feather barb was performed using analytical techniques. The morphological structure and physical properties of peacock feather barbs indicate that barbs are natural protein fibers with hollow structure. FTIR studies confirmed the protein structure, similar to chicken feather. X-ray diffraction studies revealed that the barb is a semi-crystalline protein fibre. The fibre has a β-sheet keratin structure with hexagonal unit cell. Thermogravimetric analysis results show that the barb is thermally stable up to 150°C. The barb characterization results of the studies may help the biologists in their respective field for better interpretation of research data.

